# Prime-InDel version 1: A web tool providing InDel markers for targeted regions narrowing

**DOI:** 10.64898/2026.01.29.702688

**Authors:** Yuanbao Lei, Kyaw Myo Thu, Ziqi Zhou, Linyun Xu, Chunchao Wang, Tongyao Liu, Lei Meng, Dechen Yang, Muhiuddin Faruquee, Dapu Liu, Yanbi Zhao, Xiuqin Zhao, Haizhu Chen, Zhikang Li, Wensheng Wang, Jianlong Xu, Tianqing Zheng

## Abstract

While high-density SNP genotyping enables genome-wide assays in rice (*Oryza sativa* L.) and other crops, PCR-based markers—particularly those derived from insertion–deletion (InDel) variations—remain crucial for fine-mapping. Currently, readily available primer information for high-density InDel rice markers is still limited.

We present the first version of PrimeInDel (https://rfgb.rmbreeding.cn/search/variation/primeIndel), an online tool integrating three sets of InDel markers: an 8K-set and a 316K-set from the 3K Rice Genomes (3K-RG) project, and a 22K-set compiled from published literatures. In user cases, primers from PrimeInDel proved highly effective, narrowing a cold-tolerance QTL (*qSR2*) from 2.7 Mb to 200 kb and a heading-date QTL (*qDeh1*.*1*) from 239.5 kb to 83.6 kb. Our work provides the community with a robust, PCR-based InDel primer, accessible via an online data set, which will facilitate germplasm genotyping and gene identification in rice breeding.

## Introduction

Despite the rising popularity of genome sequencing, molecular markers remain vital for breeding and genetic analysis especially those based on major-effect region targeting (Emon 2025; Dhakal et al. 2025). PCR-based markers, in particular, offer accessibility for targeted region analysis (Duan et al. 2026). Rice (*Oryza sativa* L.), a staple crop and model species, has extensive genomic resources including the 3,000 Rice Genomes (3K-RG) (Wang et al. 2018). While related databases host millions of InDel polymorphisms (Wang et al. 2020), primer information for InDel remains scattered across publications (Table 1).

**Table 1.**
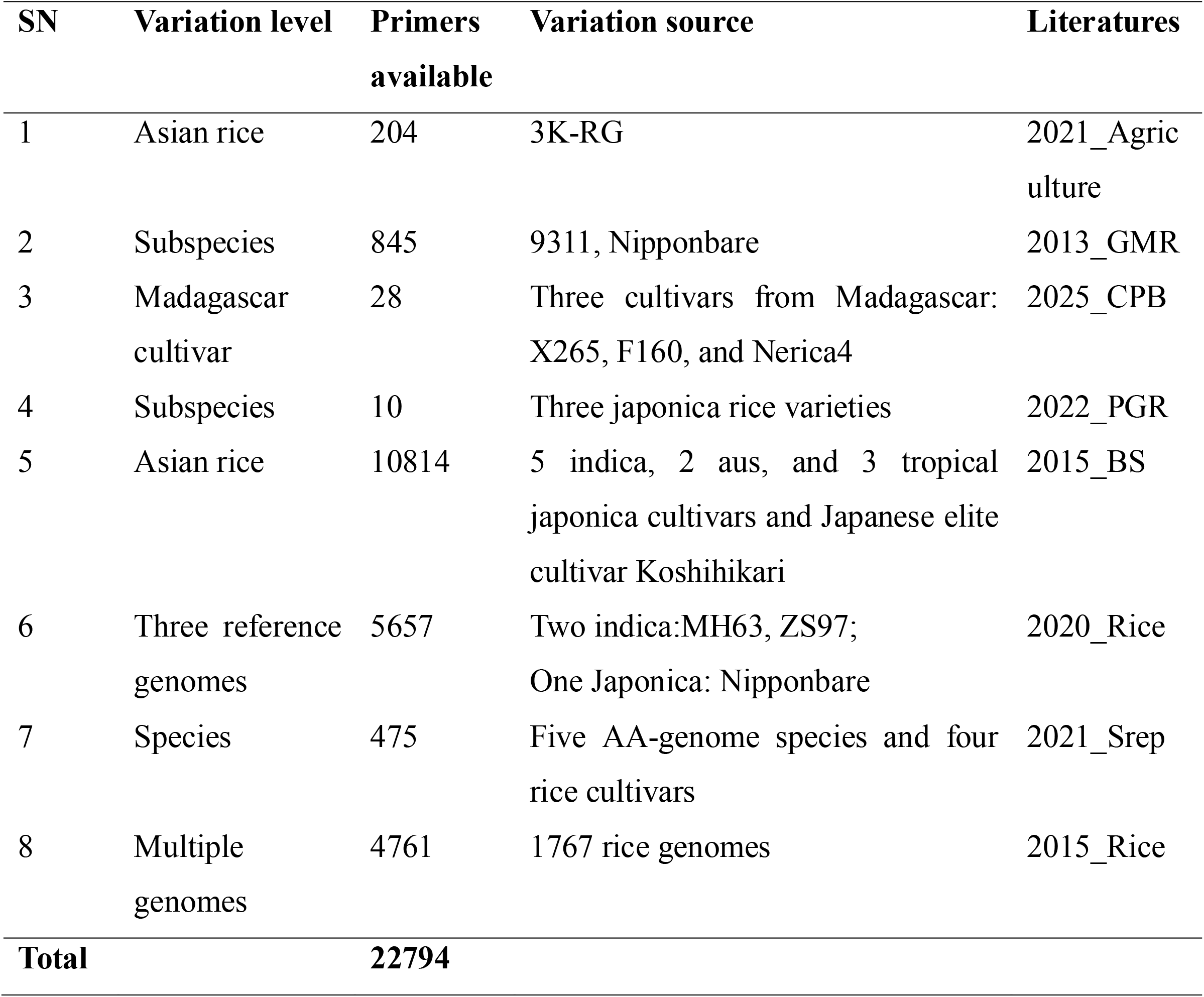
Overview of InDel marker sets available for rice from literatures.

| **SN** | **Variation level** | **Primers**<br>**available** | **Variation source** | **Literatures** |
| --- | --- | --- | --- | --- |
| 1 | Asian rice | 204 | 3K-RG | 2021_Agric<br>ulture |
| 2 | Subspecies | 845 | 9311, Nipponbare | 2013_GMR |
| 3 | Madagascar<br>cultivar | 28 | Three cultivars from Madagascar:<br>X265, F160, and Nerica4 | 2025_CPB |
| 4 | Subspecies | 10 | Three japonica rice varieties | 2022_PGR |
| 5 | Asian rice | 10814 | 5 indica, 2 aus, and 3 tropical<br>japonica cultivars and Japanese elite<br>cultivar Koshihikari | 2015_BS |
| 6 | Three reference<br>genomes | 5657 | Two indica:MH63, ZS97;<br>One Japonica: Nipponbare | 2020_Rice |
| 7 | Species | 475 | Five AA-genome species and four<br>rice cultivars | 2021_Srep |
| 8 | Multiple<br>genomes | 4761 | 1767 rice genomes | 2015_Rice |
| **Total** |  | **22794** |  |  |

Here, we introduce the first version of Prime-InDel, a one-stop online tool integrating three sets of ready-to-use InDel primers: a 22K-Ref set compiling published InDel primers, and two newly released sets—an 8K-set and a 316K-set— designed from the 3K-RG’s core and full collections, respectively. User cases for these marker sets are presented.

## Materials and Methods

### Plant Materials

For validation using a subset of the 8K-set, eight accessions from 3K-RG (Nipponbare, Akihikari, HEI-BIAO, ZHEN-XIAN 232, Kongyu131(KY131), Tonghe 886, Tonghe 899, Lianyu3252) were used.

For the QTL fine-mapping case study using the 8K-set, DNA bulks for cold tolerance from an F□population derived from cultivars Zhongnonggeng 18 (ZNG18, with Heilongjiang release No. of 2023L0088) and Zhongnonggeng 11 (ZNG11, with Heilongjiang release No. of 2024L0028) were adopted.

For fine-mapping the heading-date locus *qDeh1*.*1* (Zhou 2024), a total of 1,157 individuals from an F□RIL population from a cross involving the restorer, Minghui63 (MH63) and 7XY16, a sister line of DEH229 (Faruquee et al. 2021), a dominant early-heading line derived from MH63, were used.

### Data Source

Clean reads were obtained from the Rice Functional Genomics and Breeding (RFGB) database of the 3K-RG project (Wang et al. 2020). Sequences were aligned to the Nipponbare reference genome (IRGSP v1.0) (Kawahara et al. 2013) using BWA, and variant calling was performed with GATK. Two InDel sets were identified: 1,838,931 loci from the 454 mini-core (1.8M InDel) and 98,560,511 loci from the entire 3K-RG (98 M InDel). Unless otherwise specified, all parameters were left at their default values.

### Primer Release

Two new marker sets were designed and released: an 8K-set (8,082 markers) from the 1.8M mini-core InDels, and a 316K-set (333,460 markers) from the 98 M full-set InDels. To enhance utility in northern early *Geng*/*Japonica* rice, markers from the 8K-set were also validated using 450 sequenced accessions from Heilongjiang, resulting in a 2K subset (2,274 markers), which were then submitted to example assay.

Primers from 8K-set and 316K-set were designed in batch using BatchPrimer3 (You et al. 2008) based on ∼200 bp flanking each InDel, selecting only those with a Q-score >70 and GC content <60%.

Primers for InDel markers obtained from published literature constituted a 22K-Ref set. Among these, markers whose genomic positions were unknown in the Nipponbare genome (IRGSP v1.0) were re-aligned using BLAST (Altschul et al. 1990). The maximum-distance threshold between primer pairs for each marker was set to 2 kb.

### Web Tool Setup

The first version of Prime-InDel web tool was developed using MySQL and JavaScript, served on Nginx. Users can search for primer information in any genomic region based on physical position of reference genome (IRGSP v1.0).

### DNA Extraction and Variation Detection

Genomic DNA was extracted using the TPS method. PCR was performed with 2× M5 HiPer Plus Taq HiFi mix under standard conditions, and products were separated on polyacrylamide gels. All experimental procedures followed the standard protocols routinely employed in the Rice Molecular Breeding (RMB) lab (Faruquee et al. 2021)

### Genetic Mapping

QTL fine-mapping was performed using QTL IciMapping software (LOD >2.5). The cold-tolerance locus was delineated using 40 extreme F□individuals, and the heading-date locus was mapped using RIL population.

## Results

### The Prime-InDel Integrated Tool

We developed an online tool Prime-InDel, allowing users to search, filter, and download InDel marker information—including primer sequences—based on genomic position, dataset, or product size (Figure 1). It integrates three sets of InDel markers: an ultra-high-density 316K-set from the full 3K-RG, an 8K-set from the core collection, and a compiled 22K-Ref set from literature.

**Figure 1.**
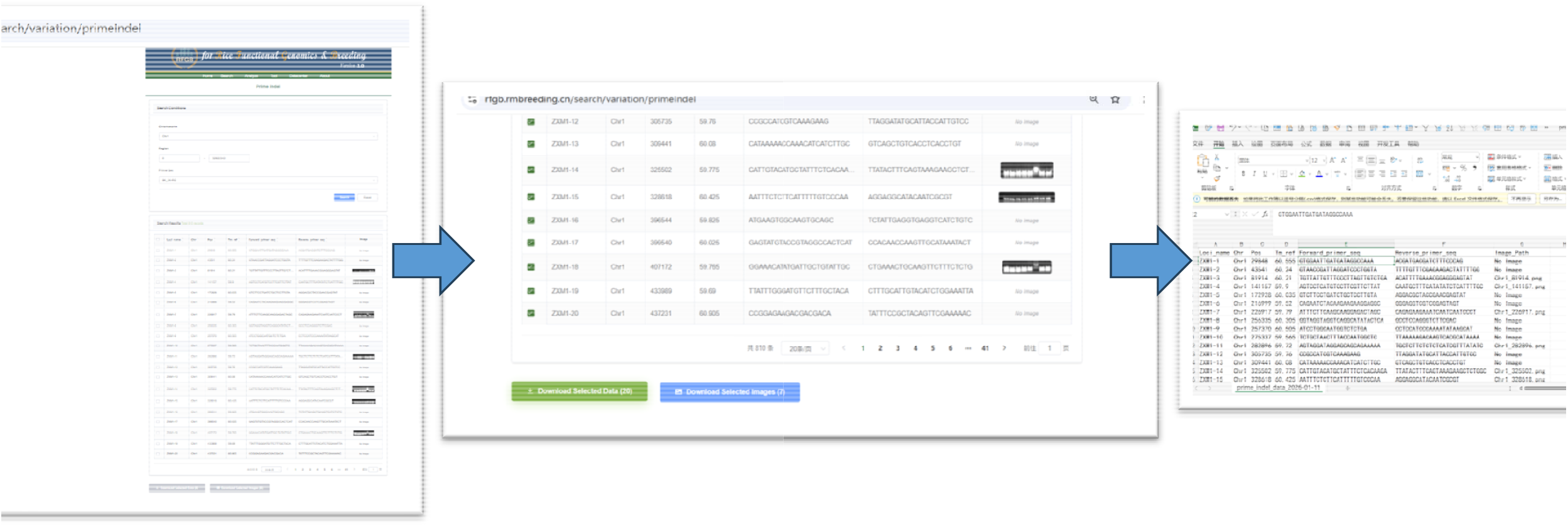
Retrieving InDel primer information by Prime-InDel.

### Amplification Efficacy of the 8K-set

As a representative of the 8K-set, a 2K subset was also assayed in eight accessions (Nipponbare, Akihikari, HEI-BIAO, ZHEN-XIAN 232, KY131, TH886, TH899, LY3252). Over 85% of markers produced clear major band, exhibiting polymorphism among the lines, demonstrating its practical value for genetic analysis and molecular breeding in this rice type. Sample gel pictures are available in the Prime-InDel (Figure 1).

### Narrowing Down QTL Regions

#### A linkage mapping-derived target region

A heading-date QTL (*qDeh1*.*1*) was previously mapped to a 239.5 kb interval (Zhou 2024). Screening by physical position in the Prime-InDel yielded 11, 17, and 52 markers from the 8K, 22K-Ref, and 316K sets, respectively. Among these, 2, 3, and 9 markers were polymorphic between parents (18.2%, 17.6%, and 17.3% polymorphism rates, Table 2, Supplementary Table 1). Using these polymorphic markers, the region of *qDeh1*.*1* was then narrowed down to about 83.6 kb with 12 candidate genes (Figure 2).

**Table 2.** InDel marker sets available for fine-mapping qDeh1.1.

| **Marker Set** | **Total in Interval** | **Polymorphic Markers** | **Polymorphism Rate** |
| --- | --- | --- | --- |
| 8K | 11 | 2 | 18.2% |
| 22K-Ref | 17 | 3 | 17.6% |
| 316K | 52 | 9 | 17.3% |

**Figure 2.**
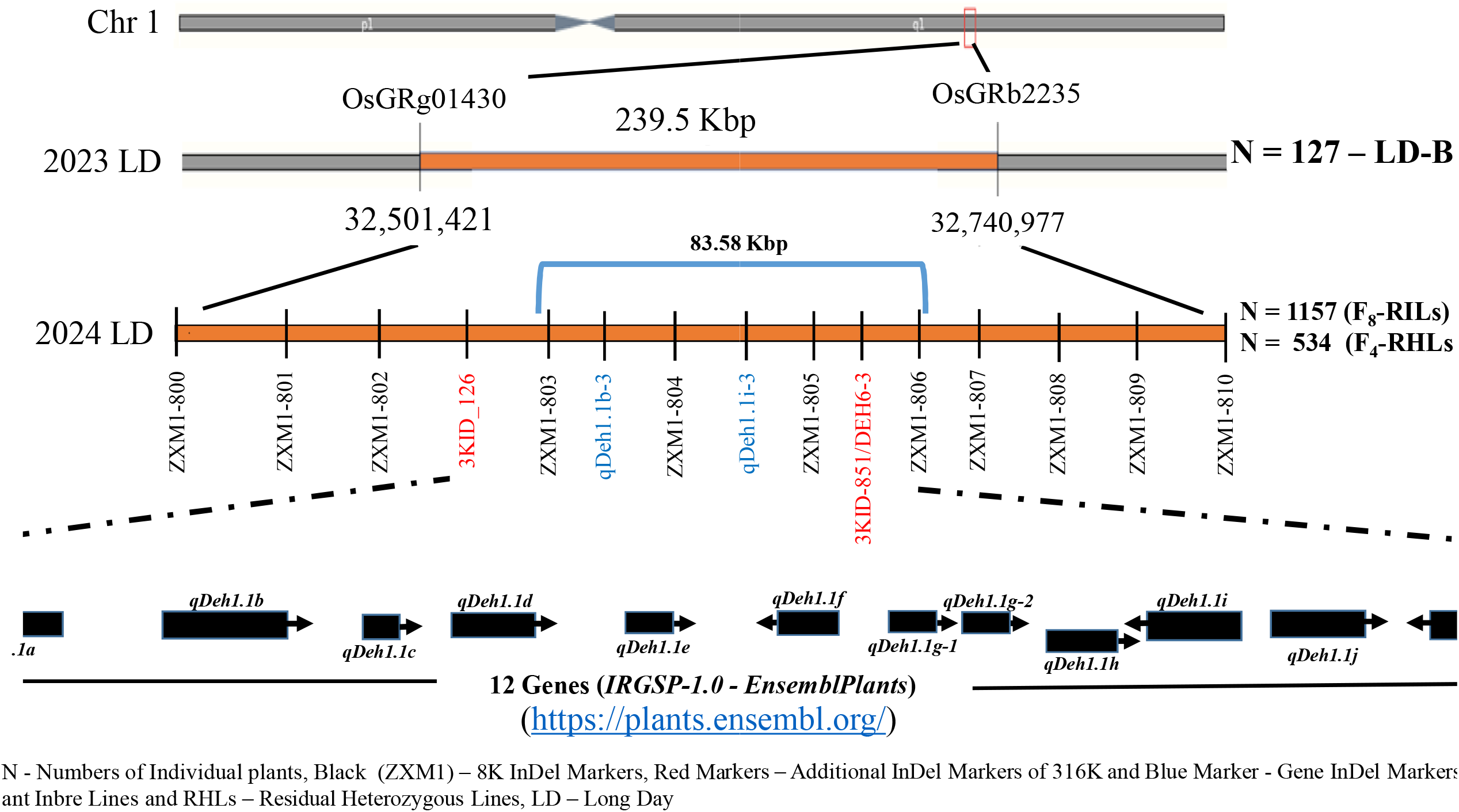
Narrowing region of *qDeh1*.*1* from linkage mapping.

#### A bulk segregating analysis (BSA)-derived target region

A previously identified cold-tolerance QTL (*qRS2*) was initially mapped on chromosome 2 (Zhang et al. 2024). Using five polymorphic markers from the 8K-set, linkage analysis with 40 extreme individuals refined the region from 2.7 Mb to about 200 kb with only 8 candidate genes (Figure 3).

**Figure 3.**
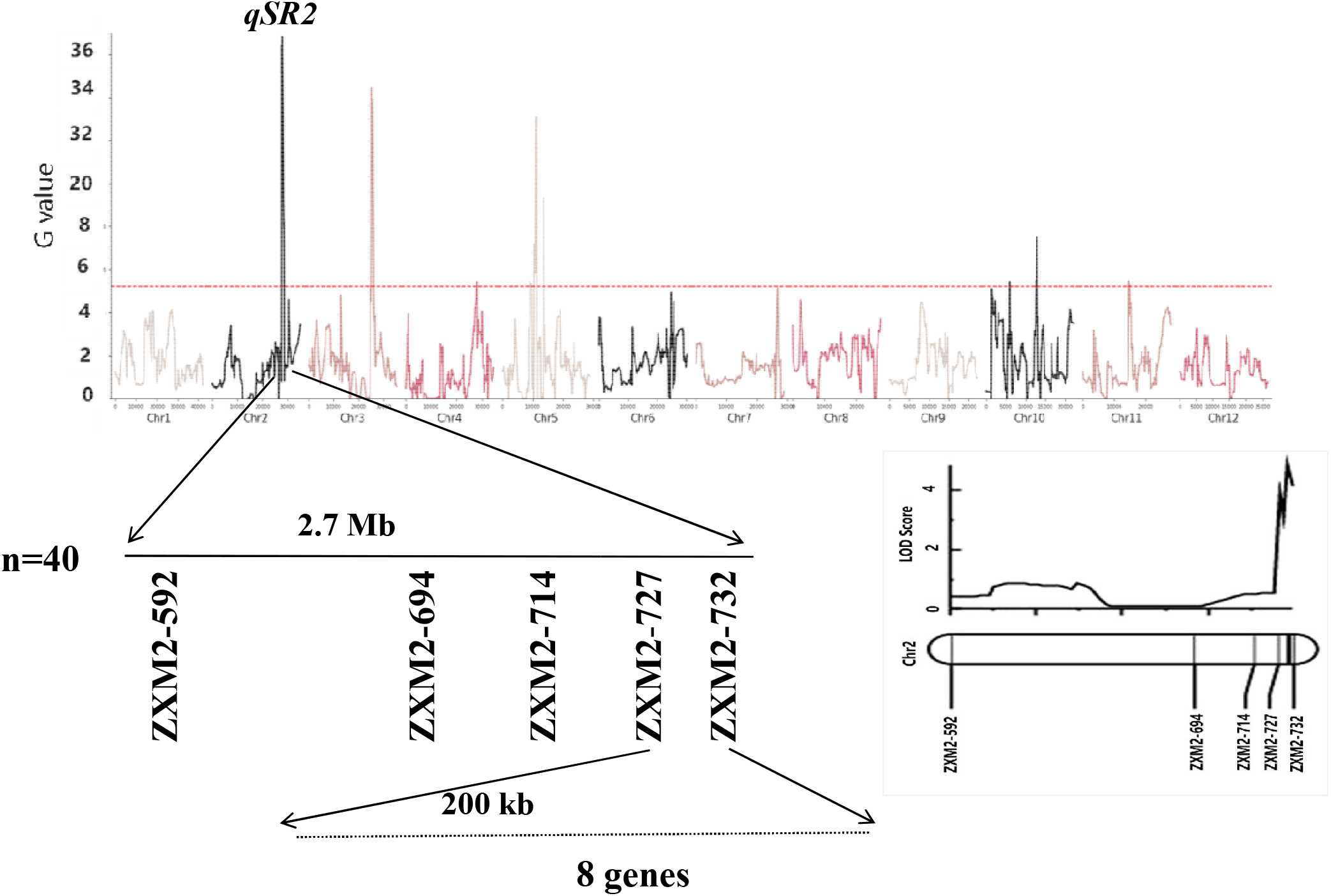
Narrowing region of *qSR2* from bulk segregating analysis (BSA)

### Data Availability

All marker data, including the 8K-set, 316K-set from 3K-RG, and the 22K-set from literatures, are available through the first version of Prime-InDel web tool (https://rfgb.rmbreeding.cn/search/variation/primeIndel).

## Discussion

This study developed and released two new genome-wide InDel primer sets (8K and 316K) derived from the 3K-RG data source and one 22K-Ref set from published literatures, along with the first version of integrated tool Prime-InDel. Our results demonstrate the advantages of these markers for polymorphism screening and QTL fine-mapping. Compared to SNPs, InDels generally exhibit satisfying polymorphism and are easier to score via routine PCR and electrophoresis, as validated in our germplasm assay.

The successful application of Prime-InDel lies in its efficient assistance in refining QTL intervals. By screening markers from the 316K-set based on physical position, we narrowed a cold-tolerance locus from 2.7 Mb to 200 kb and a heading-date QTL from 239.5 kb to 83.6 kb. This targeted strategy provides an option for researchers to accelerate fine-mapping with relatively lower costs.

Beyond fine-mapping, these marker sets hold promise for multiple breeding applications, including foreground/background selection in marker-assisted breeding, pedigree analysis, and targeted allele introgression. The online tool significantly enhances accessibility, allowing researchers to retrieve ultra-high density InDel primer information tailored to specific genomic regions or experimental needs.

In summary, this work provides the rice research community with ready-to-use InDel marker primer resources with ultra-high density, bridging the gap between high-density genomics and practical, PCR-based genotyping. As molecular breeding increasingly integrates high-resolution genetic tools, these 3K-RG-derived InDel sets and the integrated 22K-Ref set offer a scalable, cost-effective option for advancing gene discovery and cultivar development in rice and potentially other related crops.

## Conclusion

We developed Prime-InDel, an online tool integrating genome-wide InDel marker sets: an 8K-set and a 316K-set from the 3K-RG project, and a 22K-set from published literatures. The newly released 8K and 316K sets demonstrated high polymorphism and utility in germplasm screening, background selection, and QTL region narrowing. The accompanying online database provides easy access to primer information, supporting molecular breeding efforts in rice and other crops.

